# Residual force enhancement decreases when scaling from the single muscle fibre to joint level in humans

**DOI:** 10.1101/2024.06.10.598225

**Authors:** Avery Hinks, Kaitlyn B. E. Jacob, Makenna A. Patterson, Benjamin E. Dalton, Geoffrey A. Power

## Abstract

Residual force enhancement (rFE), defined as increased isometric force following active lengthening compared to a fixed-end isometric contraction at the same muscle length and level of activation, is present across all scales of muscle. While rFE is always present at the cellular level, often rFE ‘non-responders’ are observed during joint-level voluntary contractions. We compared rFE between the joint level and single fibre level (vastus lateralis biopsies) in 16 young males. *In-vivo* voluntary knee-extensor rFE was measured by comparing steady-state isometric torque between a stretch-hold (maximal activation at 150°, stretch to 70°, hold) and a fixed-end isometric contraction, with ultrasonographic recording of vastus lateralis fascicle length (FL). Fixed-end contractions were performed at 67.5°, 70°, 72.5°, and 75°; the joint angle that most closely matched FL of the stretch-hold contraction’s isometric steady-state was used to calculate rFE. The starting and ending FLs of the stretch-hold contraction were expressed as % optimal FL, determined via torque-angle relationship. In single fibre experiments, the starting and ending fibre lengths were matched relative to optimal length determined from *in-vivo* testing, yielding an average sarcomere excursion of ∼2.2-3.4µm. There was a greater magnitude of rFE at the single fibre (∼20%) than joint level (∼5%) (*P*=0.004), with ‘non-responders’ only observed at the joint level. By comparing rFE across scales within the same participants, we show the development of the rFE non-responder phenomenon is upstream of rFE’s cellular mechanisms, with rFE only lost rather than gained when scaling from single fibres to the joint level.

## 1. Introduction

Following active muscle lengthening, steady-state isometric force production is increased compared to a fixed-end isometric contraction at the same muscle length and level of activation^1^. This increase in isometric force following a ‘stretch-hold’ contraction is called residual force enhancement (rFE), and is an intrinsic property of muscle unexplained by the cross-bridge theory^2^. The mechanisms contributing to rFE originate at the sub-cellular level ^3,4^, and correspondingly, rFE has been observed in the half sarcomere ^5^, sarcomere ^6,7^, myofibril ^6^, and the single muscle fibre ^8^ and joint level in humans ^9–11^.

While rFE is always observed in single muscle fibres from human and animal preparations ^5–8,12^, rFE is not always observed *in vivo* during voluntary contractions, with some participants producing up to 34% rFE in the knee extensors, but others producing no rFE ^13–17^. This phenomenon of voluntary rFE ‘non-responders’ appears to be at least partly driven by neural factors ^17^, as spinal excitability ^18,19^ has been shown to be reduced in the rFE state, and non-responders are eliminated during electrically evoked contractions *in vivo* ^16,20^. This non-responder phenomenon raises the question of how rFE scales from the single fibre to the joint level in the same participants; however, this comparison has yet to be directly investigated.

Although rFE is characterized by enhanced force in the isometric steady-state following a stretch-hold contraction compared to a fixed-end isometric contraction at the ‘same muscle length’ ^2^, current *in-vivo* assessments of rFE are limited in that only joint angle is matched in the isometric steady state, not muscle fascicle length ^10,15,16,20^. Assessing isometric force between a stretch-hold and fixed-end contraction at the same joint angle may not necessarily equate to assessing the same muscle fascicle length due to inter-individual variability in the in-series compliance of the muscle-tendon-unit ^10,21^. Therefore, to translate *in-vivo* joint-level rFE experiments to the single fibre level, one could match fascicle length, rather than joint angle, between the isometric steady-states of the stretch-hold and fixed-end contractions. Ultrasonographic recording of fascicle length during rFE experiments ^13,22–24^, in conjunction with muscle biopsies, provides the opportunity to gain a better understanding of how rFE at the single muscle fibre level scales to joint-level voluntary contractions in humans.

The purpose of this study was to assess the scaling of *in-vitro* single fibre rFE to *in-vivo* voluntary rFE within the same participants. To achieve this, we obtained ultrasonographic recordings of fascicle length expressed as a percentage of fascicle length at the optimal joint angle (FLo) during *in-vivo* stretch-hold and fixed-end contractions, then translated the length parameters to single fibres as a percentage of optimal sarcomere length (SLo) – thereby matching experimental stretch-hold parameters across muscle scales. We hypothesized that single fibre rFE and joint-level rFE would be moderately related, with greater magnitudes of rFE at the single fibre level, and a loss of rFE when scaling to *in-vivo* voluntary contractions, contributing to the rFE non-responder phenomenon.

## 2. Methods

### 2.1. Participants

Sixteen healthy recreationally active adult males (23.3 ± 2.9 years, 80.7 ± 10.9 Kg, 180.3 ± 6.9 cm) were recruited. Participants were free of any neuromuscular disorders or lower extremity injuries and were instructed to avoid intense exercise 24 hours prior to the *in-vivo* testing visit or the muscle biopsy. Written informed consent was given prior to the study and the participants were informed of all the procedures. All procedures in this study were approved by the Research Ethics Board of the University of Guelph (REB: 22-06-009).

### 2.2. Experimental Set-Up

All participants were fitted to the HUMAC NORM dynamometer system (CSMi Medical Solutions, Stroughton, MA) with the lateral condyle of the left femur aligned with the axis of rotation. The hip was set to 100°, and the knee angle ranged from 150° to 70° (angle between the posterior aspects of the upper and lower leg; 180° = full extension) (Figure 1). Torso movement was limited with a four-point seatbelt harness, and the thigh was immobilized with an inelastic strap over the distal femur. The attachment for knee extension was secured distally on the shin via an inelastic strap. A shinpad was placed over the shin to minimize discomfort when pushing against the attachment. Two surface electromyographic (EMG) Ag-AgCl electrodes (1.5 x 1 cm; Kendall, Mansfield, MA) were placed over the muscle bellies of the rectus femoris (agonist) and biceps femoris (antagonist) with ∼4-cm interelectrode distances. A ground electrode was placed over the patella. All corresponding areas were shaved of hair and cleaned with alcohol prior to electrode placement. For evoked twitch contractions of the knee extensors, two custom-made transcutaneous stimulation pads (Aluminum foil, paper towel, tape) were coated in conducting gel and placed over the proximal (cathode) and distal (anode) ends of the quadriceps (Figure 1). The stimulator (DS7AH) was set to 400 V and a pulse width of 1000 μs. An ultrasound probe was fixed over the vastus lateralis as described in the *Ultrasound* section below. Torque, position, and stimulus trigger data were sampled at 1000 Hz using a 12-bit analog-to-digital converter (PowerLab System 16/35; ADInstruments, Bella Vista, NSW, Australia). EMG data were sampled at 2000 Hz and band-pass filtered between 10 and 1000 Hz. All data were analyzed using LabChart software (version 8; ADInstruments, Dunedin, New Zealand).

**Figure 1:**
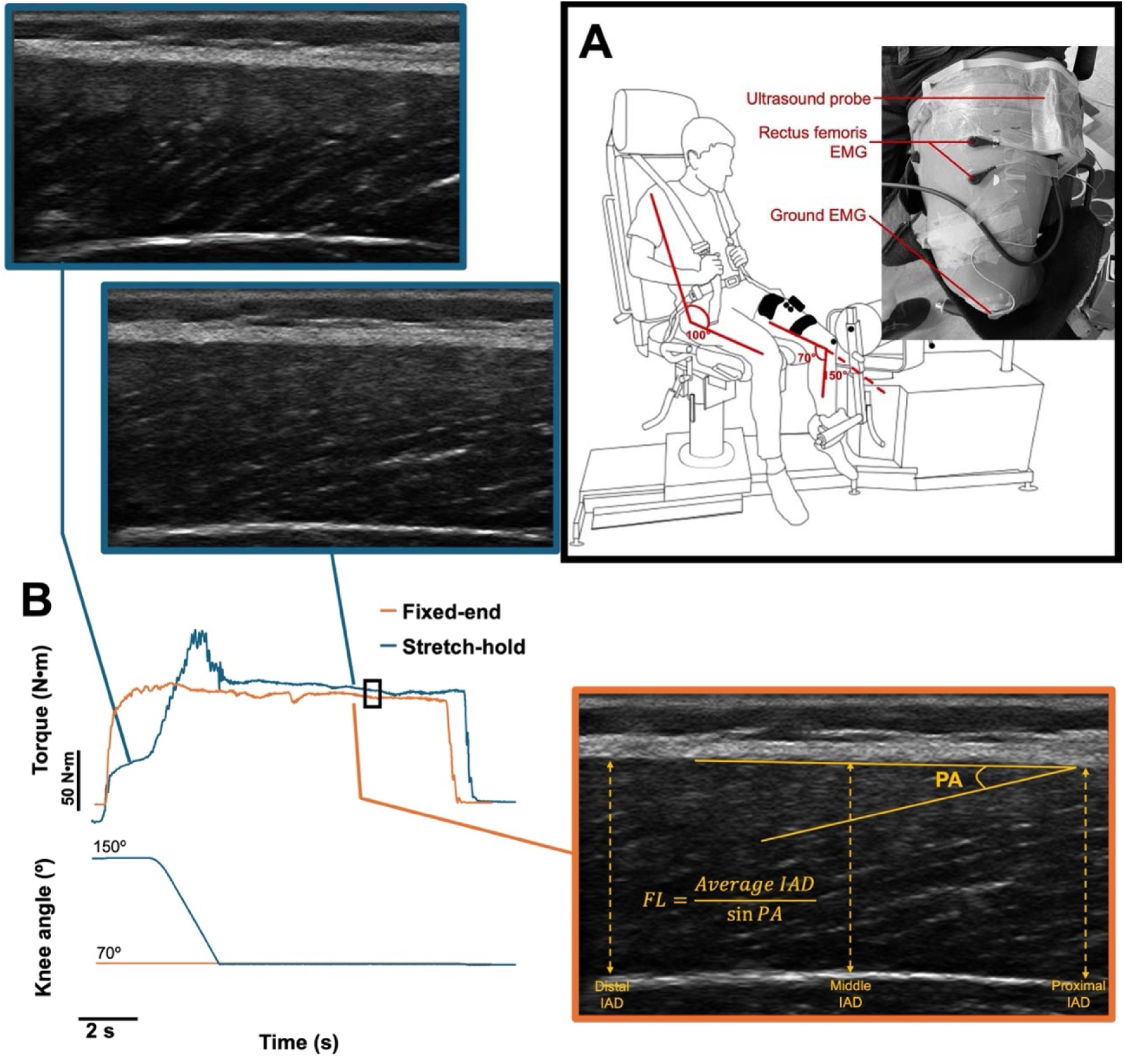
**A.** Experimental set-up. Participants were seated on the HUMAC NORM dynamometer with a hip angle of 100° and a knee angle range of 70° - 150°. Inelastic straps were fastened over the shoulders, waist, thigh, and ankle. Surface electromyography (EMG) was recorded from the rectus femoris and biceps femoris. The ultrasound probe was positioned over the vastus lateralis to record fascicle length. **B.** Torque and position traces during the fixed-end and stretch-hold, black contractions. The black box indicates the 500 ms window from which residual force enhancement was calculated. Ultrasound imaging was recorded from the left vastus lateralis in vivo. Note that the steady state isometric torque in the stretch-hold contraction is greater than that of the fixed-end contraction. Also shown are representative ultrasound images prior to stretch and during the isometric steady state of the stretch-hold contraction (upper left images) and during the isometric steady state of the fixed-end contraction (bottom right image). Fascicle length was calculated by dividing the average aeroneurosis distance (IAD) by the sine function of the pennation angle (PA).

### 2.3. Maximum Voluntary Contractions and Assessment of Voluntary Activation

A constant-current high-voltage stimulator (model DS7AH; Digitimer, Welyn Garden City, UK) was used to establish the maximal twitch torque amplitude of the knee extensors. At a knee angle of 70°, twitches were evoked by increasing the current in 20-mA increments (starting at 20 mA) until the twitch torque plateaued. The current that produced the maximal twitch torque was used for the subsequent interpolated twitch technique. After an initial maximal voluntary contraction (MVC) with no stimulations, the interpolated twitch technique ^25^ was used to confirm >90% voluntary activation of the knee extensors during MVCs at a knee angle of 70°. Peak torque generated from a superimposed twitch during the plateau of the MVC was compared to the peak of a corresponding control twitch evoked within 2 s after relaxation. The level of voluntary activation was calculated as: **Voluntary activation (%) = [1 – (superimposed twitch** / control twitch)] × 100%.

Following confirmation of ≥90% voluntary activation, additional 3-s MVCs were performed (5 MVCs in total) to prime the muscle-tendon-unit prior to collection of ultrasound images ^26^. All MVCs were separated by 3 minutes of rest. Participants were instructed to perform knee extension contractions, like kicking a ball, “as hard and fast as possible,” with visual feedback displayed on a computer monitor as a real-time torque trace. A torque guideline was provided on the computer monitor along with strong verbal encouragement throughout each MVC.

### 2.4. Muscle Ultrasound

To identify the site for ultrasound measurements, a line was drawn from the greater trochanter to the lateral condyle of the left femur using indelible ink, followed by a transverse line marking 50% of the distance between those landmarks ^22,27,28^. With the participant seated in the dynamometer and the leg fully extended, a 65-mm wide Telemed linear array probe (LV8-4L65S-3; Frequency: 8 MHz; Image depth: 60 mm) was placed just proximal to the 50% line on the lateral aspect of the vastus lateralis (Figure 1), then the probe position that produced the clearest view of muscle fascicles with the superficial and deep aponeuroses parallel in the image was determined. The probe was fixed in this location using a custom-made polyethylene probe-holder and adhesive tape (Figure 1). Transmission gel (Stevens Multi-Purpose Ultrasound Gel, Medium Viscosity) was used to improve acoustic contact and minimize transducer pressure on the skin, and care was taken to minimize pressure when taping down the polyethylene device.

The obtained video files were stored on a personal computer and specific frames were analyzed in image analysis software (ImageJ version 1.53). For measurement of fascicle length at the optimal knee angle (assumed to be optimal fascicle length; FLo), images were extracted from the videos at a time corresponding to the plateau of the MVC torque. For the 10-s rFE and isometric contractions (described below), images were extracted from the videos at the time window corresponding to the 500-ms window from which the steady-state isometric torque values were obtained. For the stretch-hold contractions, images were also extracted at the time of contraction onset (i.e., at a knee angle of 150°) to calculate the fascicle excursion **(fascicle excursion = final fascicle length – starting fascicle length)** and the start and end ranges of the active lengthening excursion as a percent of FLo (**[start or end fascicle length / FLo] × 100%**). Measurements were performed twice on two images (i.e., 4 measurements total) from each extraction by the same investigator (A.H.). Vastus lateralis fascicle lengths were determined in accordance with previous studies ^27,29,30^: the angle between the fascicle and superficial aponeurosis was defined as pennation angle, and fascicle length was calculated in each image as the inter-aponeurosis distance divided by the sine component of the pennation angle in each image (Figure 1). Here, the averaged inter-aponeurosis distance from the proximal, central, and distal ends of the image was used in calculations ^31^. Fascicle lengths were averaged between the 4 measurements obtained, and those values are reported.

### 2.5. Joint Level Data Collection

Joint-level data collection proceeded in the order of *Protocol i* to *iii*.

#### **Protocol i:** Determination of Optimal Knee Angle of Torque Production

To determine the optimal knee angle for knee extension (approximately corresponding to FLo of the vastus lateralis ^29,32^), participants performed 3-s MVCs at knee angles of 80°, 90°, 100°, 110°, and 120°. The angle corresponding to the highest torque within a 500-ms window about the MVC’s peak was deemed the optimal angle. Ultrasound was recorded throughout all contractions and for the joint angle that produced the highest torque, images were analyzed for determination of FLo. Three minutes of rest was provided between each contraction to minimize muscle fatigue. For each participant, fascicle start and end lengths during stretch-hold contractions in *Protocol ii* were expressed as a percent of FLo so that the same percentages of SLo could be used for the starting and ending sarcomere lengths in the single fibre experiments. Additionally, the fascicle lengthening velocity was normalized to FLo (units of FLo/s) so that the same velocity (units of SLo/s) during active lengthening could be used in the single fibre experiments.

#### **Protocol ii**: Maximal Voluntary Fixed-End Isometric Contractions

Ten-second fixed-end isometric MVCs were performed at knee angles of 67.5°, 70°, 72.5°, and 75°. Torque from the fixed-end contraction that most closely matched the fascicle length of the subsequent stretch-hold contraction (*Protocol iii)* was used to calculate rFE. The order of the four fixed-end isometric contractions was randomized, and they were conducted before the stretch-hold contraction to reinforce that the higher torque values observed during the stretch-hold contractions could be attributed to rFE and not the development of muscle fatigue (Paquin and Power, 2018). Five minutes of rest were provided between all trials, and strong verbal encouragement and visual feedback were provided throughout each contraction.

#### **Protocol ii**: Maximal Voluntary Stretch-Hold Contraction

The stretch-hold contractions consisted of three phases: (1) a 1-s ramp up to MVC at 150°, (2) a 2-s isokinetic lengthening at 40°/s, (3) followed by a 7-second isometric MVC at 70°. rFE was subsequently calculated in two ways: 1) comparing the stretch-hold contraction to the fixed-end contraction at 70° (i.e., joint-matching); and 2) comparing the stretch-hold contraction to the fixed-end contraction (one of 67.5°, 70°, 72.5°, or 75°) that most closely matched the stretch-hold contraction’s fascicle length (i.e., fascicle-matching). Joint-matching represents the common method of determining rFE, while fascicle-matching was necessary to replicate the *in-vivo* fascicle excursions in the single fibre experiments. Five minutes of rest were provided between all trials and strong verbal encouragement and visual feedback were provided throughout each contraction.

### 2.6. Biopsies for Single Muscle Fibre Preparation

While all 16 participants completed the joint level data collection, three dropped out prior to the biopsy sampling. Thus, biopsies were collected from n = 13. Muscle fibres were biopsied from the vastus lateralis, centered between the lateral epicondyle and the greater trochanter of the femur as described in Pinnel et al. ^8^. The fibre bundles were placed in a silicone elastomer-plated petri dish with dissecting solution. Muscle fibre bundles were then dissected approximately 0.5-1 mm in width and 3 mm in length, transferred to a tube containing 2.5 mL of chilled skinning solution, and maintained on ice for 30 minutes for permeabilization. All bundles were gently agitated to ensure equal permeabilization of all fibres in the skinning solution, with all the bundles submerged at all times. The bundles were then washed with fresh, chilled, dissection solution and gently agitated to remove any remaining skinning solution. These bundles were then stored in a separate tube containing storage solution and incubated for 24 hours at 4°C. Fresh storage solution was prepared in 0.6 mL tubes and the bundles were placed separately in each tube at -80°C, until mechanical testing began ^33^. 9 to 20 (13.46 ± 4.23) fibres were tested from each participant. Upon inspection of the individual force traces and removal of any fibres with signs of ripping or eccentric contraction-induced damage, 5 to 19 (12 ± 5) fibres from each participant were included in the final dataset. For the reporting of all single fibre data, values were averaged among all fibres within a participant to obtain a single participant-specific value. We performed this averaging to better translate the experiment back to the *in-vivo* joint level, wherein the forces produced would be “averaged” across all active fibres within a muscle.

### 2.7. Solutions for single fibre testing

The dissecting solution was composed of the following (in mM): KLproprionate (250), Imidazole (40), EGTA (10), MgCl_2_·6H_2_O (4), Na_2_H_2_ATP (2), H_2_O. The storage solution was composed of KLproprionate (250), Imidazole (40), EGTA (10), MgCl_2_·6H_2_O (4), Na_2_H_2_ATP (2), glycerol (50% of total volume after transfer to 50:50 dissecting: glycerol solution), as well as leupeptin (Sigma) protease inhibitors. The skinning solution with Brij 58 was composed of KLpropionate (250), Imidazole (40), EGTA (10), MgCl_2_·6H_2_O (4), 1 g of Brij 58 (0.5% w/v). The relaxing solution: Imidazole (59.4), K.MSA (86), Ca(MSA)_2_ (0.13), Mg(MSA)_2_ (10.8), K_3_EGTA (5.5), KH_2_PO_4_ (1), H_2_O, Leupeptin (0.05), Na_2_ATP (5.1), as well as leupeptin (Sigma) protease inhibitors. The preLactivating solution: KPr (185), MOPS (20), Mg(CH_3_COOH)_2_ (2.5), ATP (2.5). The activating solutions were composed of the same ingredients in various amounts depending on how much Ca^2+^ was needed in each solution. The composition of pCa 4.5 was as follows (mM): The activating solution (pCa 4.5) consisted of Ca^2+^ (15.11), Mg (6.93), EGTA (15), MOPS (80), ATP (5), CP (15), K (43.27), Na (13.09), and H_2_O. All solutions were adjusted to a pH of 7.0 with the appropriate acid (HCl) or base (KOH). The composition of solutions was determined by calculating the equilibrium concentration of ligands and ions based on published affinity constants (Fabiato and Fabiato, 1979). 250 units/ml of creatine phosphokinase was used in each activating solution.

### 2.8. Single Fibre Preparation

On the day of mechanical testing, single fibres were dissected in relaxing solution and tied with nylon suture knots to pins mounted between a length controller (322C, Aurora Scientific, Toronto, ON, Canada) and force transducer (403A, Aurora Scientific). Throughout all single-fibre experiments, average sarcomere length was recorded using a high-speed camera (Aurora Scientific Inc., HVSL 901A). A fitness contraction was performed with sarcomere length (SL) set to 2.8 μm. Contraction was initiated by first transferring the fibre from relaxing solution to a washing solution and then an activating solution (pCa = 4.5). Following the fitness contraction, SL was measured again and fibre length readjusted as needed.

### 2.9. Single-Fibre Residual Force Enhancement

Optimal sarcomere length (SLo) was assumed to be 2.7 µm for the human vastus lateralis ^34^. The average participant’s starting *in-vivo* fascicle length for the stretch-hold contraction was 80.7 ± 4.9% of FLo, thus an average starting single fibre SL of ∼2.18 µm was used for the single fibre stretch-hold contractions. The average *in-vivo* ending fascicle length for the stretch-hold contraction was 125.0 ± 5.4% of FLo, thus an average ending SL of ∼3.38 µm was used for the single fibre stretch-hold contractions, and for the single fibre fixed-end contractions (Figure 2). While these values stated here were the average across all participants, the actual values used were individualized to a participant’s *in-vivo* fascicle data.

**Figure 2:**
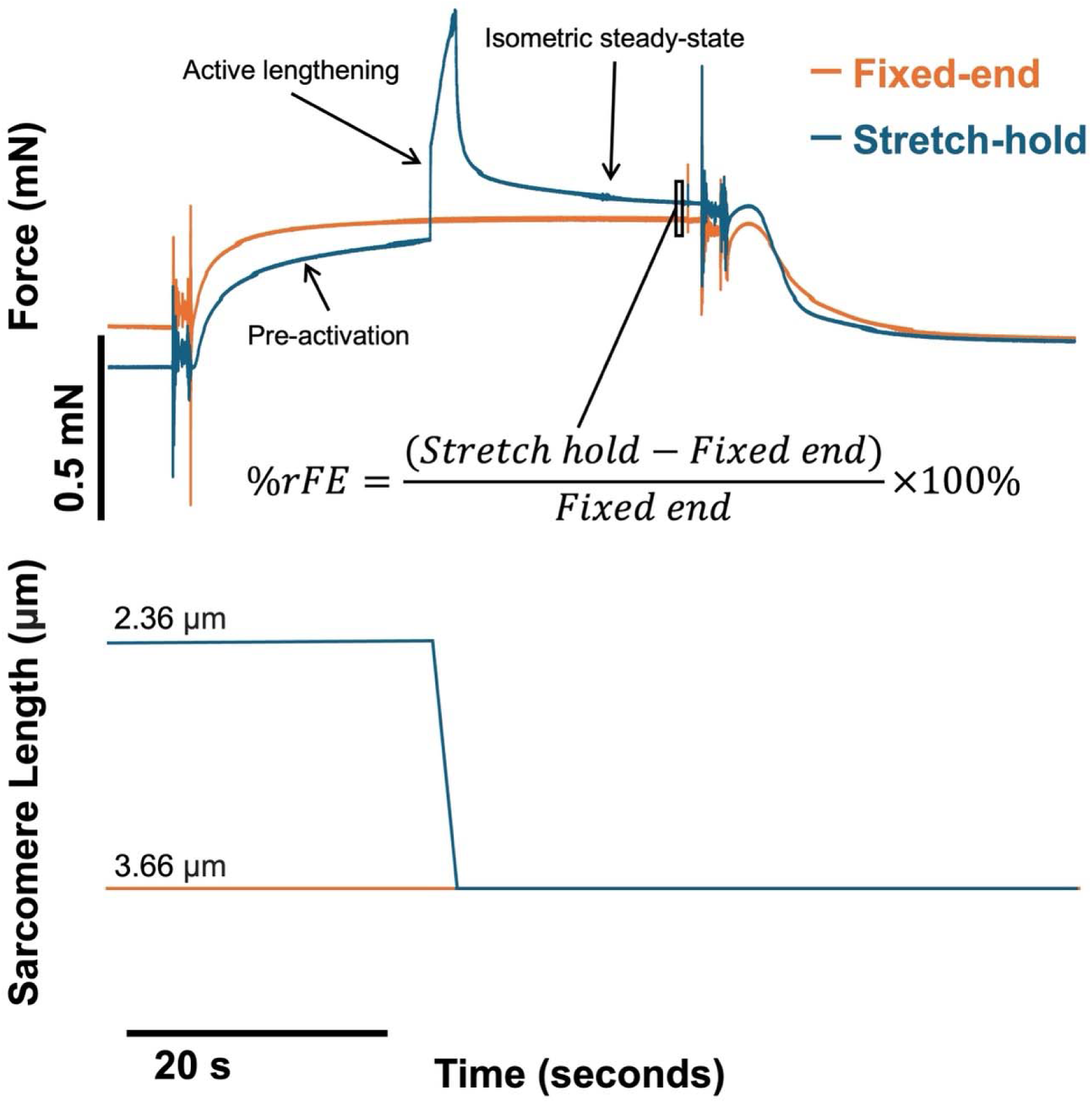
Representative data traces for single fibre force and length for a fixed-end contraction (orange) and a stretch-hold contraction (blue). Note that the steady state isometric force in the stretch-hold contraction is greater than that of the fixed-end contraction, demonstrating residual force enhancement (rFE) following active lengthening. The box indicates the 500 ms window from which rFE was calculated.

Prior to the stretch-hold contraction, fibres performed a fixed-end isometric contraction at their determined end SL. To do this, while still in relaxing solution, fibres were lengthened to their pre-determined end length, passively shortened to their starting length for 5 s, then passively lengthened back to their end length and held for 10 s before being transferred to activating solution and allowed to develop force for 40 s (Figure 2).

Following 30 s of rest, a stretch-hold contraction was performed. Similar to the fixed-end contraction, fibres were stretched to their end length before being passively shortened to their starting length. Fibres remained at this shortened length and were moved to activating solution. After 20 s of activation, fibres were actively lengthened to their end length and held at this length for 15 s before deactivating (Figure 2). The velocity during active lengthening corresponded to the fascicle lengthening velocity in optimal lengths per second determined from the *in-vivo* experiments (0.22 ± 0.02 FLo/s).

### 2.10. Data and statistical analyses

For calculation of rFE, mean torque was taken from a 500-ms window of the torque trace that was the most stable and representative of the steady-state between the stretch-hold and corresponding fixed-end isometric contraction ^20^. After the dissipation of torque transients following active lengthening, a state is reached which is deemed the “steady-state” in which torque remains mostly stable over the time ^15^. For the *in-vivo exper*iments, absolute rFE was calculated as the increase in steady-state isometric torque (Nm) in the stretch-hold contraction compared to the corresponding fixed-end isometric contraction. The same calculation was used for the single fibre experiments but in units of force (mN). For relative rFE, the absolute rFE values were represented as the percent increase (%) from the fixed-end contraction. Torque and EMG (*in-vivo* data), and force (single fibre data) were taken from the same 500-ms window following the onset of activation between the fixed-end and stretch-hold contractions, respectively.

Two-tailed, paired t-tests compared torque or force from the fixed-end isometric contractions to torque or force from the stretch-hold contractions to confirm the presence of rFE. These comparisons were performed for both the total group of participants (n = 16) and on only participants that were rFE “responders” (n = 9). A “responder” was defined as someone exhibiting greater than 1% rFE. Two-tailed paired t-tests were performed to compare rFE calculated from the commonly used joint-matching method to rFE calculated from our novel fascicle-matching method. For the 13 participants who completed both the joint-level experiments and the muscle biopsy, two-tailed paired t-tests were used to compare *in-vivo* fascicle-matched rFE to single fibre rFE. Linear regression analyses were also performed to assess the relationship between *in-vivo* fascicle-matched rFE and single fibre rFE. Where significance was detected (*P* < 0.05), effect sizes are reported as Cohen’s d. Data are reported as mean ± standard deviation.

## 3. Results

### 3.1. Voluntary activation, MVC torque, optimal angle, and FLo

For all MVCs, voluntary activation was greater than 90% (95.1 ± 3.1%), with an MVC torque of 181.9 ± 59.0 Nm. The optimal angle of torque production was 96.3 ± 6.2°, with an FLo of 98.2 ± 18.5 mm.

### 3.2. Fascicle-matched rFE: all participants

While steady-state torque in the stretch-hold contraction (149.9 ± 39.3 Nm) was on average greater than in the fixed-end contraction (146.5 ± 32.6), while there was a trend, we did not observe statistically significant rFE across the whole participant group (*P* = 0.07) (Figure 3A). The average rFE magnitude was low (relative rFE: 2.93 ± 3.89 %; absolute rFE: 3.51 ± 6.29 Nm). The stretch-hold and fixed-end contractions did not differ in agonist (stretch-hold: 0.83 ± 40 mV; fixed-end: 0.80 ± 0.43 mV; *P* = 0.27) or antagonist RMS EMG (stretch-hold: 0.16 ± 0.08 mV; fixed-end: 0.16 ± 0.08 mV; *P* = 0.82).

**Figure 3:**
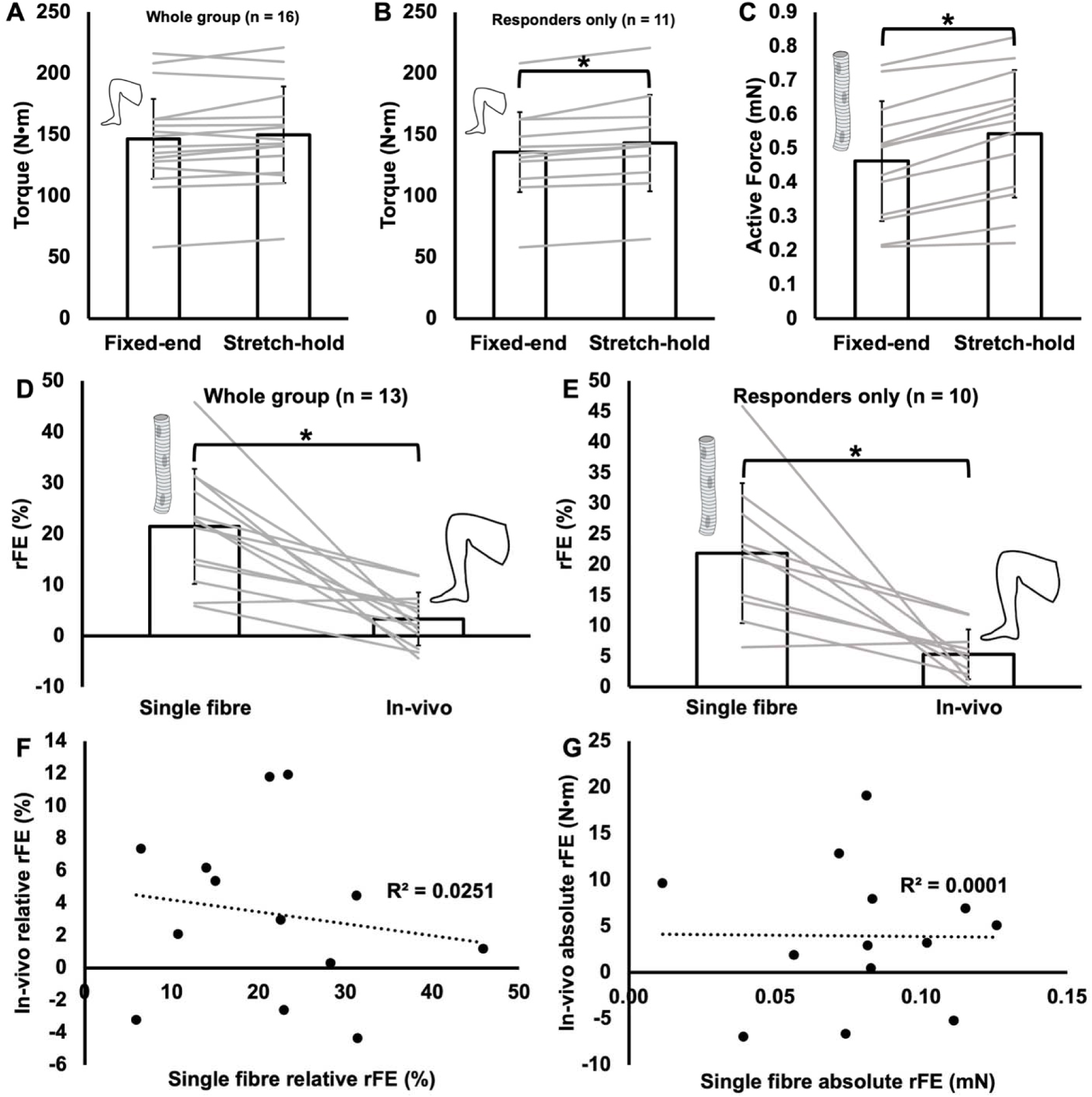
**A-B:** *In-vivo* steady-state isometric torque in the fixed-end and stretch-hold contractions for the whole participant group (A) and the residual force enhancement (rFE) responders only (B). **B:** Single fibre active force in the fixed-end and stretch-hold contractions. **D-E:** Relative residual force enhancement (rFE) from the *in-vivo* fascicle matched method compared to single fibres from the same participants for the whole group (D) and for rFE responders only (E). **F-G:** Regressions showing the lack of relationship between *in-vivo* relative rFE from the fascicle matched method and single fibre relative rFE (F) and *in-vivo* absolute rFE and single fibre absolute rFE (G). *Difference between points (*P* < 0.05). Data are presented as mean ± standard deviation. Grey lines represent individual data matched across joint-level and single fibre experiments.

### 3.3. Fascicle-matched rFE: responders only

There were a total of five rFE non-responders, representing ∼30% of the study’s participants. When reporting on only the rFE responders (n = 11), steady-state torque of the stretch-hold and fixed-end contractions were 143.15 ± 43.85 Nm and 135.84 ±40.65 Nm, respectively, and significant rFE was observed (*P =* 0.001, *d* = 0.17; relative rFE: 5.60 ± 2.09%; absolute rFE: 7.31 ± 4.61 Nm) (Figure 3B). For these rFE responders, the stretch-hold and fixed-end contractions also did not differ in agonist (stretch-hold: 0.66 ± 0.20 mV; fixed-end: 0.64 ± 0.25 mV; *P* = 0.42) or antagonist RMS EMG (stretch-hold: 0.16 ± 0.08 mV; fixed-end: 0.17 ± 0.09 mV; *P* = 0.50).

### 3.4. Comparison between single fibre rFE and in-vivo fascicle-matched rFE

rFE was present in all single muscle fibres (*P* < 0.001, *d* = 0.44) with force being 0.08 ± 0.03 mN or 21.48 ± 11.26% greater in the stretch-hold compared to the fixed-end contraction (Figure 3C). For the 13 participants who completed the biopsy procedure, relative rFE was 542% greater at the single fibre level than *in-vivo* (*P* < 0.001, *d* = 2.07) (Figure 3D). This discrepancy across scales of muscle was slightly reduced when considering only *in-vivo* rFE responders who completed the biopsy (n = 10), however, relative rFE at the single muscle fibre level was still 308% greater than *in-vivo* (*P* = 0.004, *d* = 1.93) (Figure 3E). Furthermore, there were no significant relationships between *in-vivo* and single muscle fibre relative rFE (R^2^ = 0.025, *P* = 0.606) (Figure 3F), nor between *in-vivo* and single fibre absolute rFE (R^2^ < 0.001, *P* = 0.972) (Figure 3G).

### 3.5. Comparison between rFE calculated from the fascicle-matching and joint-matching methods

Our fascicle-matching method matched the fascicle length in the isometric steady-state of the fixed-end contraction to the fascicle length in the isometric steady-state of the stretch-hold contraction with an accuracy of 0.65 ± 1.9 mm. The joint angle for the fixed-end contraction that most closely matched the fascicle length of the stretch-hold contraction was always either 70°, 72.5° or 75° (72.03 ± 1.88°). The fascicle length of the stretch-hold contraction (122.9 ± 24.3 mm) was ∼5 mm shorter (*P* < 0.01, *d* = 0.19) than the fascicle length of the fixed-end contraction performed at 70° (127.5 ± 25.9 mm), which was the angle used for calculation of rFE in the more common joint-matching method.

When data were pooled across all participants (n = 16), there were no differences in relative (*P* = 0.32) or absolute rFE (*P* = 0.21) between the fascicle-matching and joint-matching methods of calculating rFE (Figure 4A and B). Fixed-end isometric steady-state torque also did not differ between these methods among the whole participant group (*P* = 0.31). Therefore, while the fascicle-matching method succeeded in providing an isometric contraction with a more closely matched fascicle length to the stretch-hold contraction, it did not make a difference for the magnitude of rFE within the whole group of participants.

**Figure 4:**
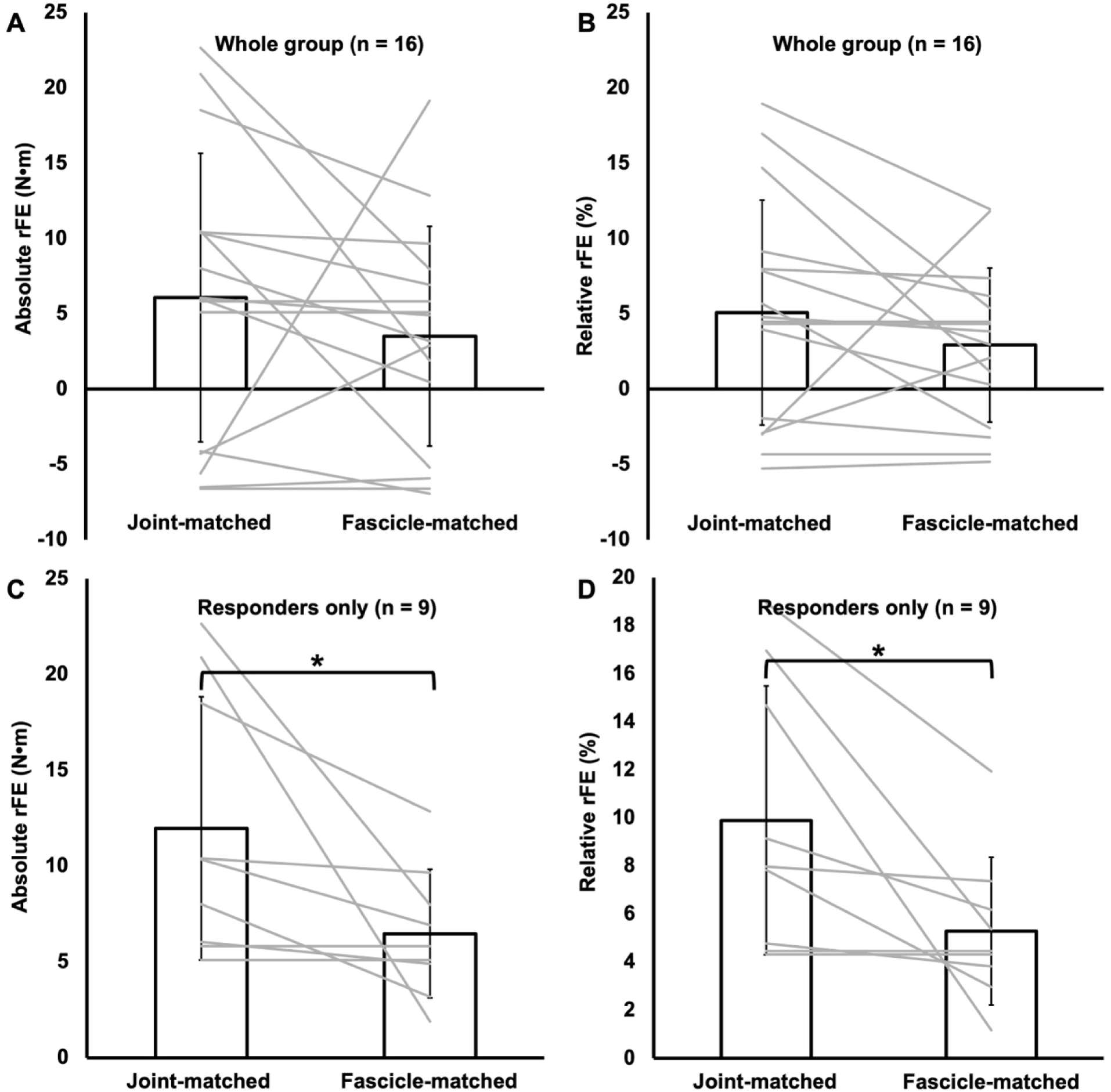
Absolute and relative residual force enhancement (rFE) for the whole group (**A-B**) and responders only (**C-D**). *Difference between points (P < 0.05). Data are presented as mean ± standard deviation. Grey lines represent individual data.

Among the fascicle-matching and joint-matching methods of calculating rFE, there were seven total rFE non-responders. When viewing data from only the rFE responders across both methods (n = 9), rFE calculated from the joint-matching method was significantly greater than rFE calculated from the fascicle-matching method (absolute rFE: *P* = 0.04, *d*= 0.81; relative rFE: *P* = 0.03*, d* = 1.2) (Figure 4C and D). Torque in the fixed-end contraction for the joint-matching method (126.82 ± 37.94 Nm) was significantly less than torque in the fixed-end contraction for the fascicle-matching method (132.44 ± 40.65 Nm) (*P* = 0.04, *d* = 0.14), which, mathematically, explains the fascicle-matching method’s lower magnitude of rFE when considering only responders in the analysis.

## 4. Discussion

The present study assessed the scaling of rFE from single muscle fibres (i.e., cellular level) to joint-level voluntary knee extension contractions, obtained by performing the same stretch magnitude as a percent of optimal length at both scales of muscle. We found that rFE was 3 to 5-fold higher in single muscle fibres than at the joint level across all participants, and non-responders were indeed only present at the joint level. These findings show that cellular level rFE is only lost, not amplified, when scaling to joint-level voluntary contractions, even when matching experimental stretch-hold parameters across muscle scales. Therefore, the non-responder phenomenon is upstream of the cellular mechanisms of rFE.

The *in-vivo* rFE values observed in the present study (1-12% for the fascicle-matching method and 4-19% for the joint-matching method) are within previously reported values of up to 34% rFE in the knee extensors for maximal voluntary contractions ^13,22,24,30,35^. Furthermore, the magnitudes of rFE we observed in single fibres (6-46%) are within previously reported ranges in single fibres from the human vastus lateralis ^8^. Our observation of ∼30% of participants being rFE non-responders for *in-vivo* voluntary contractions is also consistent with previous studies ^36–38^.

### 4.1. Tempering of rFE’s cellular mechanisms when scaling from the single fibre to the joint level

The most accepted mechanism for rFE exists at the sub-cellular level. Upon activation, titin becomes stiffer and shorter ^39^, thereby contributing more ‘passive force’ to total force production following stretch as compared with a ‘purely isometric’ contraction ^11^. The magnitude of rFE we observed in single fibres was 3 to 5-fold greater than in *in-vivo* voluntary contractions from the same participants even when matching experimental stretch-hold parameters across muscle scales (Figure 3). Furthermore, and rather surprising, there were no relationships between single fibre and *in-vivo* voluntary rFE (Figure 3F-G). By comparing rFE across these scales within the same participants, these observations highlight that no further rFE is gained beyond the sub-cellular based mechanisms driving rFE when scaling up to the joint level, contributing to the development of the non-responder phenomenon.

Certain confounding factors when scaling from the single muscle fibre to the joint level may temper rFE’s cellular based mechanisms. Our fascicle-matching method allowed us to lessen the impact of in-series compliance of the tendon as a confounding factor (by matching actual fascicle length rather than inferring fascicle length via joint angle), however, other non-contractile tissues such as intramuscular connective tissue can also buffer force transmission across the muscle-tendon unit ^40–43^. At the joint level, there are also other knee extensor muscles besides the vastus lateralis contributing to force production. Hence, while the vastus lateralis is the largest of the quadriceps muscles and is most representative of the whole knee extensor group’s torque-angle relationship ^32^, our results may not be generalizable to the other quadriceps muscles due to differences in force-length properties. For example, the vastus intermedius undergoes greater fascicle length changes during eccentric contractions than the vastus lateralis since the fascicles of the vastus intermedius insert directly onto bone ^44^. Additionally, the vastus medialis and rectus femoris tend to reach optimal angle at more extended and more flexed knee angles, respectively, than the vastus lateralis ^32^. Lastly, neural contributions to force production during *in-vivo* voluntary contractions are not present in single fibre experiments. Most notably, spinal excitability is reduced in stretch-hold compared to fixed-end isometric contractions ^45^, and this effect is likely modulated by the inhibitory Golgi tendon organ-evoked reflex, which is elevated in the force-enhanced state ^19^. Since this inhibitory reflex is activated by muscle-tendon unit stretch, the large-magnitude joint excursion (70°) that we used was likely a large contributor to the loss of rFE when scaling from the single fibre to the joint level, and the appearance of non-responders ^17^.

### 4.2. Recommendations for future research regarding fascicle-matching vs. joint-matching

When only the rFE responders (n = 9) data were included in analysis, we observed greater rFE using the joint-matching method (∼10%) compared to the fascicle-matching method (∼5%) (Figure 4). The joint-matching method’s greater rFE occurred mathematically because the fixed-end contractions used to calculate rFE for the joint-matching method had lower torque values than the fixed-end contractions used to calculate rFE for the fascicle-matching method. These lower fixed-end torque values likely occurred because, in the fascicle-matching method, some of the fixed-end contractions used to calculate rFE were at knee angles of 72.5° or 75°, which are closer to the optimal angle (∼100°) than 70°, the only knee angle used in the joint-matching method. During eccentric contractions of the vastus lateralis, much of the stretch is taken up by the tendon, limiting the amount of stretch that the contractile tissue (i.e., the fascicles) can undergo ^21,46^. Thus, it is understandable that vastus lateralis fascicle length in the isometric steady state following active lengthening would be shorter than during a fixed-end isometric contraction at the same joint angle, like in the present study. A similar result was observed by ^47^, with on average ∼2-3 mm shorter fascicle lengths measured in the rFE state compared to the isometric state for the same joint angle.

From these results, it is tempting to conclude that joint-matched rFE (i.e., the commonly used method for quantifying rFE *in vivo*) could overestimate actual rFE if accounting for fascicle length. However, we only saw a difference between the joint-matching and fascicle-matching methods when removing all participants who were non-responders in either condition. When including all participants in the comparison between joint-matched and fascicle-matched rFE, we saw no difference between the two methods. Altogether, joint-matching yields rFE results that are not significantly different from fascicle-matching within a full participant pool. For future studies investigating rFE, we recommend that researchers attempt to match fascicle length in the isometric steady-state between the fixed-end and stretch-hold contractions if rFE non-responders are to be removed from the analyses.

### 4.3. Methodological considerations

The size of our ultrasound probe did not allow visualization of fascicles from end to end in an ultrasound image, so the linear extrapolation method was used to measure fascicle length, and there is some error associated with this technique ^48^. However, our vastus lateralis fascicle length values are within previously reported ranges ^30,44,49,50^ and fascicle length measurement via the linear extrapolation method has been validated ^49^. Additionally, we assessed rFE during maximal voluntary contractions, however, rFE can also be assessed *in vivo* through submaximal electrically stimulated contractions. While including electrically stimulated rFE in the present study may have provided a closer comparison to single fibres (i.e., by removing neural contributions to force production) ^16,20^, it would have required activation of the single fibres at a submaximal pCa, which would have been difficult to implement alongside the experiments already being performed. Thus, comparison of *in-vivo* submaximal electrically stimulated rFE to submaximal single fibre rFE is an important direction for future studies. Lastly, the participants in this study were all males, so it is possible our results are not generalizable to females; however, no sex differences in rFE have been observed previously ^20^.

## 5. Conclusion

We investigated the scaling of rFE from the single fibre level to *in-vivo* voluntary contractions within the same participants by matching experimental stretch-hold parameters across muscle scales. We observed 3 to 5-fold greater rFE in single fibres than *in vivo* despite employing the same relative magnitude of stretch at both levels. By comparing rFE across these scales within the same participants, we can see clearly that the development of the rFE non-responder phenomenon is upstream of the sub-cellular mechanisms of rFE, and rFE is only lost rather than gained when scaling up to the joint level. The modifiability of rFE through resistance training has been investigated previously in the context of inducing muscle morphological adaptations ^15,51,52^. It is increasingly clear, however, that neural factors are largely responsible for the non-responder phenomenon, thus an important future direction from this research is to modulate neural factors to improve the scalability of rFE’s sub-cellular mechanisms.

## Acknowledgements

We thank Dr. Yeeshale Chetty and Dr. Erin Weersink for performing the muscle biopsies.

## Authors’ contributions

AH, KBEJ, and GAP conceived and designed the research. AH, KBEJ, MAP, and BED performed experiments. AH, KBEJ, BED, MAP, and GAP analyzed data, interpreted results, prepared figures, and drafted manuscript. AH, KBEJ, and GAP edited and revised the manuscript. All authors have read and approved the final version of the manuscript and agree with the data and order of presentation of the authors.

## Data availability

All data generated or analysed during the study are available from the corresponding author upon request.

